# IMPACT OF FLUORESCENT DYES ON MUTATIONS IN NEXT GENERATION SEQUENCING LIRBARY PREPARATION

**DOI:** 10.64898/2026.04.26.720908

**Authors:** Vincent L. Butty, Pranav Patel, Stuart S. Levine

## Abstract

DNA labelling fluorescent dyes such as ethidium bromide have long been considered to be highly mutagenic during DNA replication. While recent studies have pushed back on this narrative, the intercalative nature of these dyes continues to raise the possibility that these dyes can induce mutations. The iconPCR instrument by n6tec uses fluorescent dyes to measure amplification in real time and to adjust cycling conditions. However, since this use of qPCR is preparative and not analytical, mutations introduced by fluorescent dyes would be propagated into the sequencing reaction. To address the impact of these dyes on downstream analyses, we have performed routine mutation calling as well as mutational signature analysis on samples amplified using the iconPCR in the presence of either SYBR or EvaGreen. Sequence analysis revealed very minimal impacts of dyes on the reactions, largely within the noise regimen with only subtle changes in mutation rates seen. Mutational signature analysis was unable to identify any key signatures assignable to the dyes in either substitutions or indel domains. The mutational impact of intercalating dyes during fluorescence-guided amplification is therefore minimal and can be disregarded in all but the most sensitive NGS applications.

## Introduction

Intercalating dyes have been used to detect and measure DNA for several decades.^1^ These dyes fluoresce robustly in the context of double-stranded DNA, and the variable fluorescence led to the invention of quantitative PCR, which incorporates optical detection into the typical PCR reaction to measure the amount of fluorescence after each cycle. By measuring the cycle at which fluorescence crosses a set threshold, these instruments can robustly quantify the amount of input material included in each reaction. The intercalating mechanism of these dyes, inserting between adjacent base pairs and distorting the DNA helix, raises a direct concern regarding mutagenicity. Intercalating dyes have long been classified as suspected mutagens and possible carcinogens in laboratory safety documentation, a classification supported by early bacterial mutagenicity assays.^2^ More recent studies have found that mutagenic potential requires metabolic activation and may be substantially lower than initially feared,^3^ though the possibility that these dyes could introduce errors during DNA replication remains an open question.

The n6tec iconPCR (Individually Controlled PCR) takes this reaction a step further. Rather than applying uniform cycling conditions across an entire plate, the instrument independently monitors and controls each of 96 wells in real time, halting individual reactions as they exit the exponential amplification phase. This per-well control offers a significant advantage over conventional fixed-cycle thermocyclers in NGS library preparation, where both over- and under-amplification present critical challenges. Under-amplification yields insufficient material and may necessitate re-amplification, introducing both delay and variability. Over-amplification, particularly reactions driven into the PCR plateau phase, is associated with GC and composition bias, increased duplicate rates, chimera formation, and the generation of spurious higher-molecular-weight heteroduplex products, all of which can compromise downstream sequencing analyses.^4,5,6^

The use of intercalating dyes in the iconPCR therefore raises a specific concern: dye-induced errors introduced during preparative amplification would be propagated into the sequencing library and potentially confound downstream analyses. To assess this risk, we employed mutational signature analysis alongside conventional variant calling. Mutational signatures are characteristic combinations of mutation types arising from specific mutagenic processes; a mutagen acting through a defined mechanism, such as base intercalation, would be expected to produce a recognizable pattern of mutations attributable to that mechanism.^7^ We have therefore performed detailed analysis of reference samples prepared using the iconPCR in the presence of intercalating dyes. Bulk analysis revealed minimal changes in both single nucleotide variation rates and insertion or deletion frequencies. Mutational signature analysis revealed minimal signatures not associated with known mechanisms. Thus, amplification with the iconPCR should have minimal impact on NGS analyses except in the most sensitive applications.

## Methods

### Human genomic DNA and library preparation

Human genomic DNA from GIAB reference material NA12878 was obtained from the Coriell Institute.^17,18^ DNA was enzymatically fragmented to a target insert size of approximately 150–200 bp using Ultra II FS DNA Library Prep kit, New England Biolabs E7805. Following fragmentation, end repair, dA-tailing, and ligation of P5/P7 Illumina-compatible adapters were performed per manufacturer’s protocol. Ligation products were purified using NEBNext Sample Purification Beads (NEB E6552A), at a 0.8X ratio.

### iconPCR amplification

Amplification was performed on the n6tec iconPCR instrument using NEB Q5 High-Fidelity DNA Polymerase. Libraries were prepared in the presence of SYBR Green I, EvaGreen, or EvaGreen Plus at 1X (final concentration) alongside a no-dye control, for 10 cycles under the following conditions: initial denaturation at 98°C for 30 seconds, 10 cycles at 98°C for 10 seconds and 65°C for 75 seconds, final extension at 65°C for 5 minutes and hold at 4°C. Dye concentrations and cycle counts were intentionally set above the iconPCR’s standard stop-point parameters to provide a stringent test of mutagenic potential. Amplified libraries were purified using NEBNext Sample Purification Beads (NEB E6552A), at a 0.8X ratio.

### Sequencing, alignment and quality control

Libraries were quantified by Agilent Fragment Analyzer and by qPCR and sequenced on an Element AVITI using 2×75nt reads to a depth of approximately 100 million reads per sample (93–138M). Sequencing reads from the four conditions were aligned against GRCh38 using bwa mem (VN:0.7.17-r1188).^8^ Quality control metrics were calculated (including number of aligned reads, multiply-mapping reads, and number of unique 20-mers in the top 10M reads) and samples were checked for contamination against a collection of reference genomes.

### Variant calling

Each sample was mapped against the Genome In A Bottle (GIAB) GRCh38-based reference assembly GRCh38_GIABv3_no_alt_analysis_set_maskedGRC_decoys_MAP2K3_KMT2C_KCNJ 18.fasta, obtained from https://ftp-trace.ncbi.nlm.nih.gov/giab/ftp/release/references/, using bwa mem (v. 0.7.12-r1039, -t 16 cores).^8^ Resulting BAM files were sorted and indexed using samtools v.1.3. Variant calling was performed using bcftools mpileup (v. 1.10.2+htslib-1.10.2) with parameters -B -x -d 1000000 --threads 8 -A -O v, using the above reference assembly.^9,10^ VCF files were filtered against the GIAB benchmark VCF for sample NA12878_HG001 (HG001_GRCh38_1_22_v4.2.1_benchmark.vcf.gz, retrieved from https://ftp-trace.ncbi.nlm.nih.gov/ReferenceSamples/giab/release/NA12878_HG001/latest/GRCh3_8/) using the bcftools isec function (flags -c all -O v).^18^ No additional filtering was performed. SNV rates were calculated as total observed variant bases divided by total bases sequenced; indel rates were calculated analogously.

### Mutational signature analysis

Variant positions, reference and derived alleles from condition-specific VCF files were processed using the Signal mutational spectra elicitation webservice for single and double substitutions and indels (https://signal.mutationalsignatures.com/).^11,12^ Signature reconstruction was first performed on the absolute mutational spectra of each condition using SigProfilerAssignment (COSMIC v3.4, default activity thresholds).^14^ Dye-imparted mutational spectra were then assessed by subtracting the frequency of each variant class in the no-dye control from the corresponding frequencies in each of the three dye-treated samples. These difference frequencies were multiplied by the number of observed sequence variants of the corresponding class in each dye-treated sample, rounded, and floored to 0, then submitted as 96-, 78-, or 89-channel variation catalogues to Signal. Signature reconstruction was repeated on these different spectra using SigProfilerAssignment with the same parameters. Indel classes were assessed using a recently-published taxonomy.^13^ For base count level processing, non-reference sites were annotated using the Signal’s companion library signature.tools.lib (https://github.com/Nik-Zainal-Group/signature.tools.lib), including the indelsig.tools.lib-contained functions in the R statistical environment (v. 4.2.0). Corresponding reference and derived allele counts were retrieved from the original vcf files and tallied as above-mentioned variation catalogues for downstream processing in Signal.

## Results

To evaluate the gross impact of intercalating dyes present in the amplification reaction on NGS library quality, P5/P7-anchored libraries were prepared from enzymatically sheared human genomic DNA (GIAB reference sample NA12878) in the presence of SYBR Green I, EvaGreen, or EvaGreen Plus using NEB Q5 High-Fidelity DNA Polymerase on the n6tec iconPCR instrument (Figure 1A). Relatively high concentrations of dye and cycle counts that exceed the standard stop point of the iconPCR were selected to amplify any mutagenicity from the dyes while remaining within the performance envelope of the iconPCR. Libraries were sequenced on an Element AVITI to maximize sensitivity to any dye-induced mutations. Global library metrics showed that dye-containing reactions produced libraries of comparable quality to the no-dye control, with only subtle differences in insert size distribution (Figure 1B) and GC content (Figure 1C) observed.

**Figure 1.**
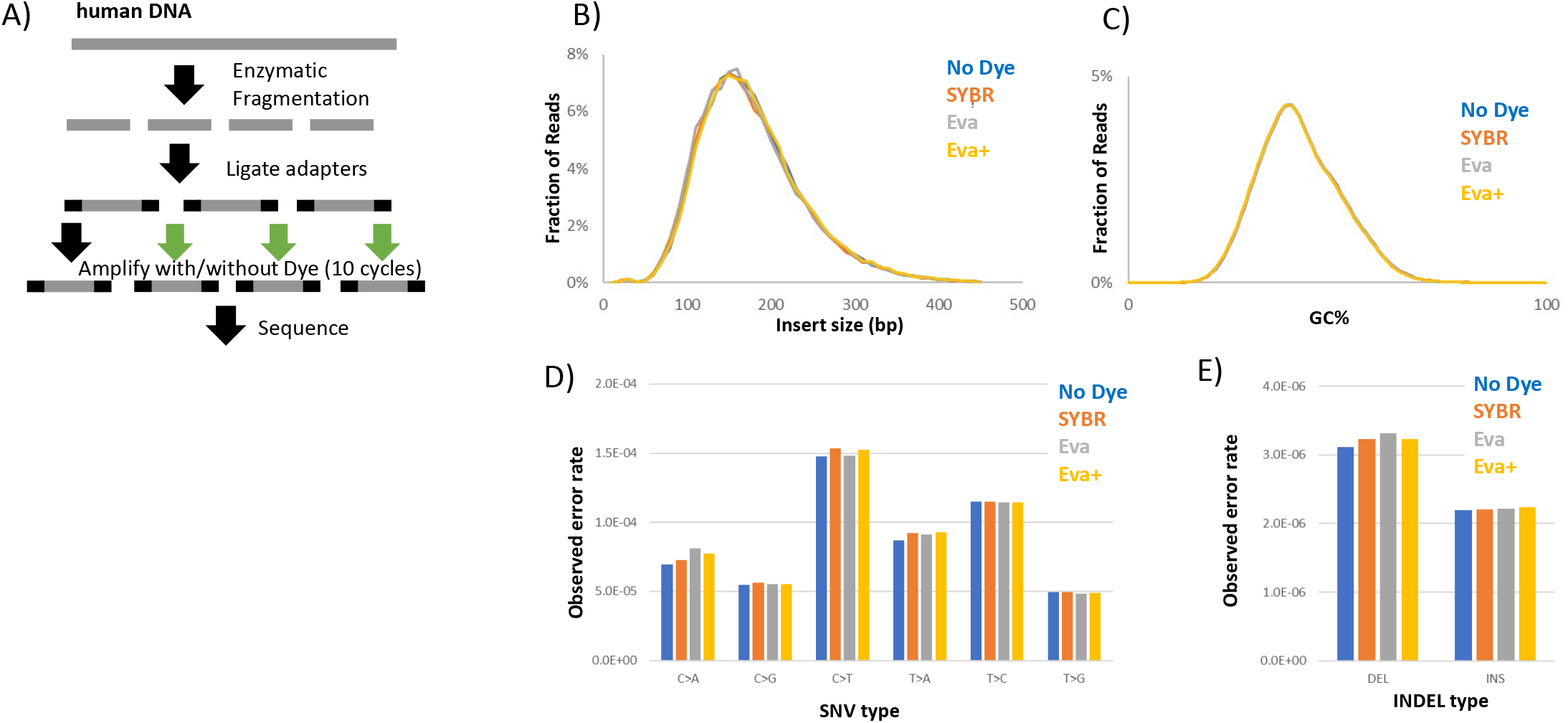
Impact of intercalating dyes on NGS library quality and mutation rates. **(A)** Schematic of the experimental workflow. **(B)** Histogram of insert lengths (bp) for libraries prepared under each dye condition. SYBR: SYBR Green I; Eva: EvaGreen; Eva+: EvaGreen Plus. **(C)** Frequency distribution of GC fraction for libraries prepared under each dye condition. **(D)** Observed single base mutational error rates by substitution type (C>A, C>G, C>T, T>A, T>C, T>G), normalized to total bases sequenced, for libraries prepared under each dye condition. **(E)** Observed indel error rates by event type (deletion, insertion), normalized to total bases sequenced, for libraries prepared under each dye condition.

The primary impact of intercalating dyes is expected to be in the induction of mutations. Single base mutational rates were characterized relative to the total number of bases sequenced (since all errors are expected to be technical in origin). Mutation analysis found very similar error rates among all the samples, regardless of the presence of dyes, with dye-containing samples increasing the error rate by less than 3.5% above the no-dye control, with SYBR Green I showing very slightly lower substitution error rates than either EvaGreen formulation. Base-by-base substitution errors showed similar rates of transitions and transversions (Figure 1D). Indel mutation rates were also only slightly perturbed by the presence of the dyes with ∼5% more deletions and 1–2% additional insertions observed, but still extremely close to background rates (Figure 1E).

We next investigated changes in mutational signatures caused by the presence of dyes in the amplification reaction. Mutational signatures are more sensitive than SNP analysis in that they include the context of the nucleotide change and are frequently highly correlated with the mechanism of action.^7^ Thus, a mutation that is associated with base intercalation would be expected to have a specific signature which could help separate it from the background. Since all observed variants are expected to be technical errors in this reference sample design, any signature enriched in dye-containing conditions would be attributable to the dyes themselves.

Mutational spectra across all dye conditions were superficially indistinguishable from the no-dye control (Figure 2A). Attempted decomposition of these spectra into known SBS signatures using SigProfilerAssignment revealed no dominant signature attributable to dye exposure; since this reaction system is entirely in vitro, a genuine dye-induced mutagen would be expected to produce a simple and recognizable signature. Reconstruction from known SBS types confirmed no systematic difference between conditions. To increase sensitivity, the no-dye spectrum was subtracted from each dye-containing condition to isolate any residual dye-specific signal (Figure 2B). These difference spectra showed only modest increases, predominantly in C>T and T>C transitions across all dyes, with EvaGreen and EvaGreen Plus also showing minor C>A transversions. Attempted signature reconstruction of the difference spectra identified no dominant signal; the highest contributors were SBS4 in EvaGreen (0.54) and SBS8 in SYBR and EvaGreen Plus (0.486 and 0.421, respectively), with no consistent pattern across conditions. Thus the differential nucleotide transition signature represents a likely novel artifactual signature.

**Figure 2.**
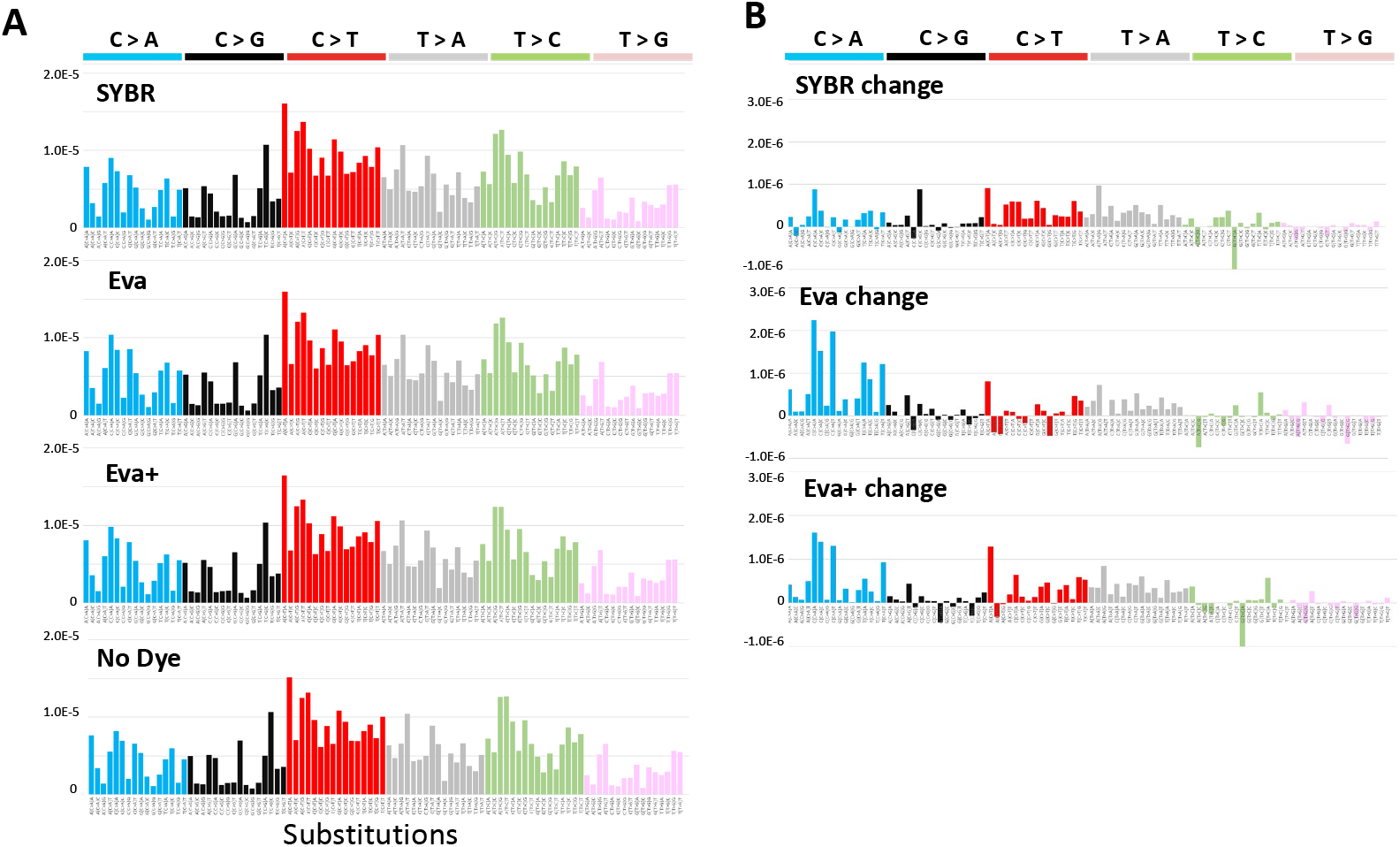
Single base substitution spectra and dye-induced changes in libraries amplified with intercalating dyes. **(A)** Observed mutational spectra across 96 trinucleotide substitution contexts, normalized to total bases sequenced, for libraries prepared under each dye condition and the no-dye control. Substitution classes (C>A, C>G, C>T, T>A, T>C, T>G) are color-coded as indicated. SYBR: SYBR Green I; Eva: EvaGreen; Eva+: EvaGreen Plus. **(B)** Dye-imparted substitution changes, calculated by subtracting the no-dye spectrum from each dye-containing condition.

We next looked for changes in indel signatures associated with dye presence in the amplification reaction (Figure 3A). Given that an intercalated dye molecule occupies a position structurally analogous to a base pair, intercalating dyes might be expected to preferentially induce single-base insertions, particularly in repetitive sequence contexts where slippage is most likely. Absolute indel spectra were highly similar across all conditions with no major shift in pattern observed. Consistent with the bulk indel rates observed in Figure 1, deletion events exceeded insertions in all dye conditions; notably, no enrichment of single-base insertion events was observed in any specific sequence context, ruling out a context-specific insertion signal masked by the overall deletion excess. Focusing on the differences between the spectra (Figure 3B) revealed modest but consistent shifts in indel patterns across dye-containing conditions. The most prominent signals were a modest increase in 2–4 bp deletions (∼4–10% above the no-dye control, with SYBR Green I being the most pronounced) and deletions of a single C preceding an A (∼4–10% above the no-dye control). A corresponding decrease in deletions of 5 bp or greater (∼2–6% below the no-dye control) was also observed. In all cases, the changes represented a minority of observed mutations.

**Figure 3.**
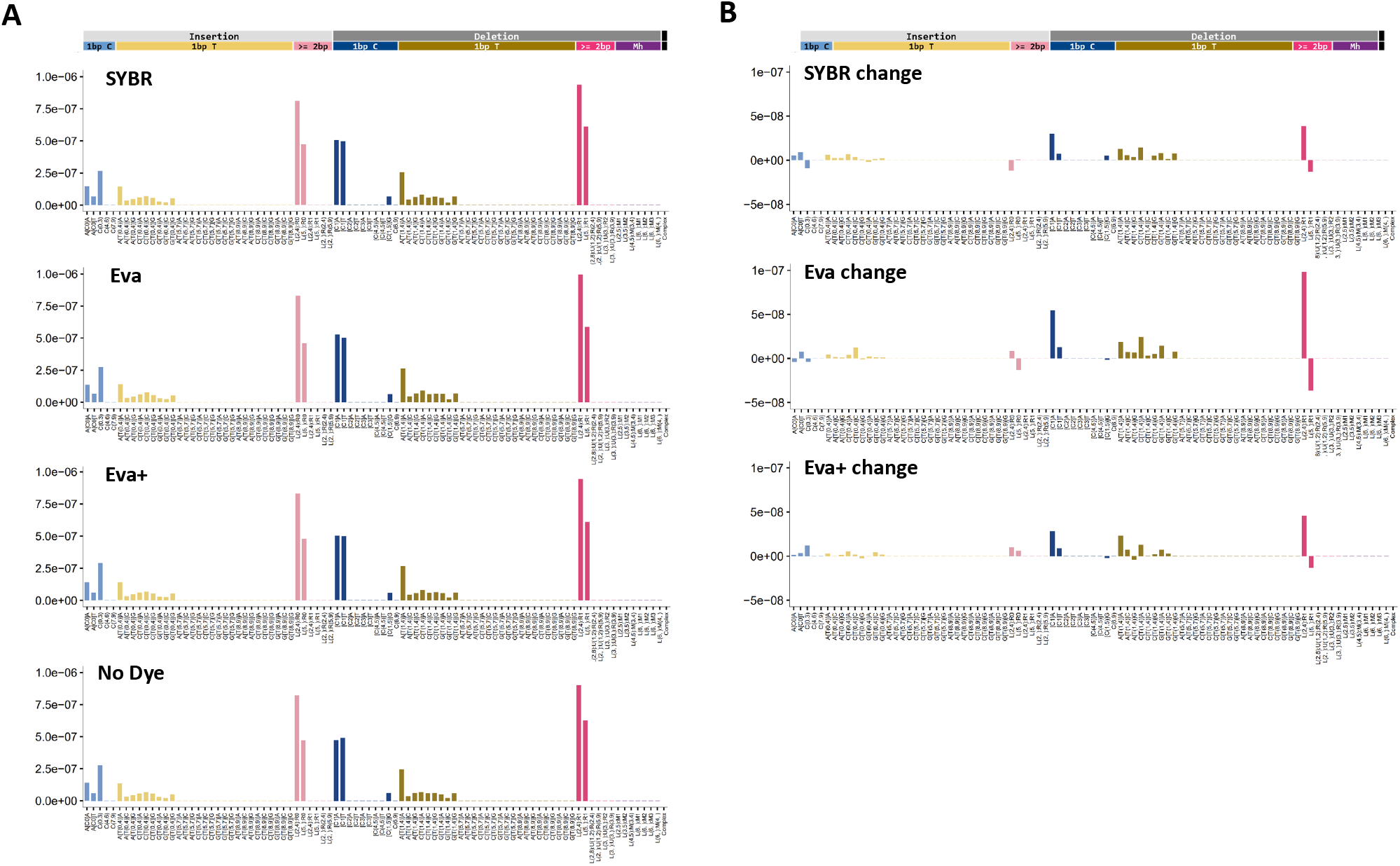
Indel spectra and dye-induced changes in libraries amplified with intercalating dyes. **(A)** Observed indel spectra across insertion and deletion contexts as defined by Koh et al.,^13^ normalized to total bases sequenced, for libraries prepared under each dye condition and the no-dye control. Complex mutations are not shown. SYBR: SYBR Green I; Eva: EvaGreen; Eva+: EvaGreen Plus. **(B)** Dye-imparted indel changes, calculated by subtracting the no-dye spectrum from each dye-containing condition.

We next examined whether the observed indel changes corresponded to any known mutational signatures. Reconstruction of the absolute indel spectra identified no dominant signature attributable to dye exposure, with the sum of all assigned signatures reaching a cosine fit of only 0.77 across conditions. Focusing on the difference spectra (Figure 3B), where only dye-induced changes should be represented, yielded even poorer overall fits (0.67–0.71). The largest contributors to these difference spectra included signatures associated with TOP1-linked deletions (InD4a) and APOBEC activity (InD9a), neither of which has a plausible mechanistic link to base intercalation. Attempted modeling against a panel of experimental chemical genotoxin signatures similarly failed to identify a coherent signal, with the highest-scoring match, 6-Nitrochrysene, explaining only a minority of the EvaGreen difference signatures, and cosine fits ranging from 0.45–0.56 across conditions. The indel difference spectrum therefore lacks any coherent signature attributable to dye exposure, consistent with the substitution analysis.

## Discussion

Intercalating dyes present during iconPCR amplification had minimal impact on mutational rates across all analyses performed. Single-nucleotide variant rates in dye-containing conditions were elevated by less than 3.5% above the background error rate of the no-dye PCR control, and indel rates were similarly close to background, with modest increases in short deletions but no enrichment of single-base insertions in any dye condition. Mutational signature analysis revealed no coherent signature attributable to dye exposure in either substitution or indel space, with recovered signals mapping to artifactual or biologically implausible signatures with no consistent pattern across conditions. These results were obtained under deliberately stringent conditions, with dye concentrations and cycle counts set above the standard iconPCR operating parameters, suggesting that the rates observed here represent an upper bound on real-world dye-induced mutagenicity.

The observed error rates are unlikely to meaningfully impact results in the large majority of NGS workflows, including bulk whole-genome and whole-exome sequencing, RNA-seq, ChIP-seq, and ATAC-seq. Applications that depend on variant detection at the lowest measurable allele frequencies warrant more careful consideration. Minimal residual disease detection requiring discrimination of somatic variants at allele frequencies as low as 0.001–0.01%,^15^ where any systematic elevation in technical error rates could contribute false-positive calls, is one such domain. Single-cell DNA sequencing for clonal analysis, where amplification errors cannot be averaged across a bulk population and are a recognized source of noise,^16^ represents a further such domain. Finally, de novo mutational signature analysis in translational or mechanistic contexts warrants particular awareness. The consistent pattern of C>T and T>C transitions in the substitution difference spectra, together with the modest enrichment of 2–4 bp deletions in the indel difference spectra, provides a recognizable fingerprint that should make any dye contribution detectable at the level of the mutational spectrum, though it cannot speak to the origin of any individual variant. Accordingly, caution is warranted in the most sensitive applications operating at the limits of detection, where even a modest shift in the error background may have meaningful consequences for results.

## Acknowledgements

The authors are thankful for Drs. John Essigmann, Robert Croy, Bogdan Fedeles, and members of the MIT BioMicro Center for their constructive comments and discussions on this manuscript. This work was funded by the National Cancer Institute of the US National Institutes of Health (NIH) under award P30-CA14051. Data from the manuscript are available at https://fairdomhub.org/studies/1454 including links to all data repositories. PP is an employee of n6 bioTec.

## Notes

https://fairdomhub.org/studies/1454

